# Macrophages Mediate Antiviral Immunity and Repair of Type 2 Alveolar Epithelial Cells in a Human Stem Cell Model

**DOI:** 10.1101/2025.04.15.648867

**Authors:** Declan L. Turner, Hannah Baric, Katelyn Patatsos, Sahel Amoozadeh, Michael See, Kathleen A. Strumila, Jack T. Murphy, Liam Gubbels, Elizabeth Ng, Andrew Elefanty, Melanie Neeland, Shivanthan Shanthikumar, Sarah L. Londrigan, Mirana Ramialison, Fernando J. Rossello, Ed Stanley, Rhiannon B. Werder

**Author notes:** Corresponding author: Rhiannon Werder.

## Abstract

The lung alveoli are constantly exposed to inhaled pathogens and inorganic hazards, relying on robust defence mechanisms to maintain homeostasis. Alveolar macrophages and type 2 alveolar epithelial cells (AT2s) collaborate to orchestrate protection. Compromised defence can dysregulate immunity and repair, leading to acute and chronic respiratory diseases. To better understand these processes and drive therapeutic discovery, human model systems that capture key cell interactions are essential. Here, we develop the first induced pluripotent stem cell (iPSC)-derived platform that integrates AT2 cells and macrophages in an air-liquid interface culture. Coculture enhanced AT2-specific gene expression and lipid synthesis, while macrophages actively phagocytosed AT2-derived surfactant. iPSC-derived AT2s supported macrophage survival by producing M-CSF and coculture promoted an alveolar macrophage-like phenotype. Additionally, during respiratory infection macrophages played a crucial role in modulating proinflammatory signalling, enhancing antiviral immunity, and restricting viral replication. Furthermore, we identify a role for iPSC-derived macrophages in epithelial repair, with VEGF signalling to macrophages increasing epithelial permeability. We present an iPSC-derived air-interface platform to study AT2-macrophage interactions in homeostasis, infection, and repair, providing insights into their potential roles in the initiation and progression of respiratory diseases.

## Introduction

The lung alveoli are constantly exposed to inhaled pathogens and pollutants, requiring robust defence mechanisms. When these defences are dysregulated, lung diseases can arise from persistent injury and abnormal repair. Alveolar macrophages act as frontline immune cells, clearing harmful particles and promoting tissue repair ^1,2^, while type 2 alveolar epithelial cells (AT2s) support immune regulation, surfactant production, and epithelial repair ^3,4^.

Respiratory infections carry the highest disease burden, exceeding major public health threats like cancer and heart disease, measured by disability-adjusted life years ^5^. Alveolar macrophages have a critical role in resolving respiratory viral infections in mice ^6–9^. AT2s are common targets for respiratory viruses and primary producers of infectious virus ^10–12^, and dysfunction and death of AT2s is associated with respiratory failure ^13–15^. Given the close proximity of alveolar macrophages and AT2s in the alveoli, studies have explored how immune responses to respiratory viral infections are influenced through cell crosstalk. For instance, AT2-derived surfactant protein (SP)A and SPD inhibit macrophage activation, in part through binding to SIRPα on alveolar macrophages ^16–19^. During influenza infection in mice, epithelial cells supress alveolar macrophage activation through CD200 ^20^, positing that CD200R is critical for maintaining alveolar macrophage homeostasis. However, the expression of these ligand-receptor pairs discovered in mice differs in the human lung (LungMAP ^21^). Thus, the study of respiratory viral infections and the discovery of new antivirals will necessitate the development of a human platform that incorporates the alveolar epithelium and macrophages.

The application of induced pluripotent stem cells (iPSCs) to create disease-relevant human cells is transforming the study of respiratory diseases. We and others have developed protocols using defined conditions to derive iPSC-derived AT2s (iAT2s) and macrophages (iMacs) ^22–25^. Crucially, these mimic *in vivo* functions, including surfactant biosynthesis ^24^ and immune function ^25,26^, and therefore can be applied to successfully model genetic ^24,27,28^, acquired ^29–31^ and infectious ^25,26^ lung diseases. Moreover, we have demonstrated that the reproducibility and scalability of these iPSC-derived platforms support drug screening ^25,32^. To model AT2-macrophage interactions with iPSCs, previous work has introduced iMacs to alveolar-organoids ^33,34^. However, because these organoids are polarized “apical-in,” they do not readily permit the study of respiratory infections and lack the native air-interface of the lungs, which we have previously shown to induce iAT2 maturation ^29^.

Here we develop a novel iPSC-derived platform, incorporating AT2s and macrophages in an air-liquid interface. We demonstrate that coculture promotes maturation and function of both iAT2s and iMacs. Following viral infections, iMacs heavily influence proinflammatory signalling and antiviral immunity which limits viral replication. Finally, we find that iMacs influence epithelial repair through VEGFA-VEGFR2 signalling. This human model system has the potential to improve disease modelling and drug screening for various acute and chronic conditions affecting the lung alveoli.

## Methods

### Human iPSC maintenance

Human iPSCs were cultured in StemFlex (ThermoFisher), mTeSR1 or TeSR-E8 (StemCell Technologies) on plates coated with Matrigel (Corning #354277). iPSC lines used in this study have been previously described: BU3 NGST CRISPRi ^31^, SCT3010 (MCRIi032-A, RRID:CVCL_D0I2; Stem Cell Technologies/MCRI), PB001 ^35^. Ethical approval for the generation and/or use of human iPSCs were obtained from Murdoch Children’s Research Institute (MCRI) and Boston University and were carried out in accordance with the National Health and Medical Research Council of Australia (NHMRC) regulations.

### Directed differentiation of iPSC-derived type 2 alveolar epithelial cells

Directed differentiation of iAT2s was performed as we have previously described ^22,24^. In brief, iPSCs were differentiated to CXCR4+ cKit+ definitive endoderm using the STEMdiff Definitive Endoderm Kit (StemCell Technologies). Cells were then dissociated with Gentle Cell dissociation reagent (StemCell Technologies), replated on growth factor reduced Matrigel (Corning #354277)-coated plates and cultured in anteriorisation media for three days (complete serum-free differentiation medium (cSFDM) base media, supplemented with 2 mM Dorsomorphin (Tocris, 3093/10) and 10 mM SB431542 (Tocris, 1614)). To induce NKX2-1+ lung progenitors, cells were moved to cSFDM supplemented with 3 mM CHIR99021 (R&D Systems, RDS442310), 10 ng/mL recombinant human BMP4 (R&D Systems, 314-BP), and 100 nM retinoic acid (Sigma, R2625). On day 14–15, cells were dissociated with 0.05% trypsin (Thermofisher) and NKX2-1+ lung progenitors were purified (based on NKX2-1-GFP, or CD47^hi^CD26^lo^) by FACS using a FACSAria Fusion (BD Biosciences). Sorted NKX2-1+ lung progenitors were embedded in growth factor reduced Matrigel (Corning #356230) droplets and supplemented with 3 mM CHIR99021, 10 ng/mL rhKGF (R&D Systems, RDS251KG050), 50 nM dexamethasone (Sigma, D4902), 0.1 mM 8-Bromoadenosine 30,50 cyclic monophosphate sodium salt (Sigma, B7880), and 0.1 mM 3-Isobutyl-1methylxanthine (IBMX; Sigma, I5879) in cSFDM (“CK-DCI media”). CK-DCI media was replaced every 2-3 days. iAT2s were serially passaged every two weeks, and re-sorted when needed based on NKX2-1-GFP+ SFTPC-tdTomato+ expression or CPM+, as described ^22,31^. After each passage, cultures initiated with single cells were supplemented with 10 mM Y-27632 (‘‘Y’’; Tocris, RDS125410) in CK-DCI media. Following for 2-3 days in Y-27632, medium was replaced with CK-DCI alone.

To create air-liquid interface (ALI) cultures, iAT2s were dissociated using 0.05% trypsin to generate a single cell suspension. 200,000 cells were plated onto growth-factor reduced Matrigel coated 6.5 mm Transwells (Costar) in CK-DCI media, as described ^22,29^. 2-3 days later the apical media was removed to create an ALI. Basolateral media was changed every 2-3 days. To create iPSC-derived type 1 alveolar epithelial cells (iAT1s), 200,000 iAT2s were plated on 6.5 mm Transwells in cSFDM supplemented with 10 mM LATS-IN-1 (MedChemExpress, HY-138489), 50 nM dexamethasone, 0.1 mM 8-Bromoadenosine 30,50 cyclic monophosphate sodium salt, and 0.1 mM IBMX (“L-DCI” media), as described ^36^. Trans-epithelial electrical resistance (TEER) measurements were taken with a Millicell ERS-2 Voltohmmeter (Millipore, MERS00002).

### Directed differentiation of iPSC-derived macrophages

Directed differentiation of iMacs was performed as we have previously described ^25^. In summary, on day 0 iPSCs were dissociated and resuspended in Magec media ^25^ supplemented with 20 ng/mL VEGF (R&D Systems, 293-VE), 20 ng/mL SCF (synthesised by CSIRO, Australia), 5 ng/mL FGF2 (R&D Systems, RDS233FB500), 10 ng/mL BMP4 (R&D systems, 314-BP), 10 uM Y-27632, 0.5 uM CHIR99021, and 10 ng/mL Activin A (R&D Systems, 338-AC). To form embryoid bodies (EBs), cells were cultured in non-tissue culture treated dishes (Greiner Bio-One, 628161) on a Ratech rotating platform at 60 rpm in a 37°C incubator, as described ^37^. On day 1 and 3, the media was changed by allowing EBs to settle without centrifugation then aspirating media and replacing with Magec media supplemented with 20 ng/mL VEGF, 20 ng/mL SCF, 10 ng/mL FGF2, and 10 ng/mL BMP4. From day 6 to 13, media was supplemented with 20 ng/mL VEGF, 20 ng/mL SCF, 10 ng/mL FGF2, and 25 ng/mL M-CSF (Peprotech, #300-25), and from day 13 onwards comprised 50 ng/mL M-CSF. After day 9 or 10, media changes were performed by centrifuging at 300 x g for 5 minutes to pellet EBs. Cells were filtered through a Falcon 40 µm cell strainer (Corning, 35240) at day 15-16 to remove EBs, iMacs were analysed by flow cytometry, then used for experiments over the next 2-3 weeks. Media was replaced twice weekly by centrifuging at 300 x g for 5 minutes to pellet cells, then replenishing media containing 50 ng/mL M-CSF.

To create a source of “definitive” iMacs, we first created iPSC-derived haematopoietic stem cells (iHSCs), as previously described ^38^. At day 14 of this protocol, iHSCs were plated in 50 ng/mL M-CSF and 5 ng/mL SCF in SPELS media ^38^ in non-tissue culture treated dishes on a Ratech rotating platform at 60 rpm in a 37°C incubator. Media was replaced every 2-3 days. After one week, media was changed to 50 ng/mL M-CSF in SPELS and cultured for another three weeks. Media was replaced twice weekly by centrifuging at 300 x g for 5 minutes to pellet cells. The cell pellet was then resuspended in fresh media and returned to the incubator.

In some experiments, iMacs were cultured in low-attachment 96 well plates in CK-DCI media, supplemented with 50 ng/mL M-CSF, or 50 ng/mL GM-CSF (Peprotech, #300-03). Alternatively, CK-DCI media conditioned by 2-3 days dwelling in 3D iAT2 cultures was added to iMacs. To block M-CSF or GM-CSF signalling, conditioned media cultures were treated with 4 ug/mL anti-M-CSF and/or anti-GM-CSF (R&D Systems, RDSMAB216SP and RDSMAB615SP).

To visualise live macrophages in cultures, iMacs were labelled with CellTrace CFSE or Violet at room temperature for 20 minutes, as per manufacturer’s instructions (ThermoFisher Scientific, C34554 and C34557).

### Coculture of iAT2 and iMacs

To establish iAT2-iMac cocultures at ALI, iAT2s were seeded in Transwells, as described above. 4-7 days post initiation of ALI, 20,000 iMacs were added to the apical surface of each Transwell resuspended in 5-10 µL CK-DCI (this volume was reabsorbed or evaporated in 2-3 days). Experiments were conducted 4-21 days post addition of iMacs, as indicated. A 10:1 ratio of iAT2:iMacs was selected based on the estimated frequency of type 2 alveolar epithelial cells and alveolar macrophages in the human lung ^39,40^.

### Flow cytometry and cell sorting

Endoderm cells were stained for CXCR4 (BD Bioscience, 555974) and c-Kit (Biolegend, 313206) and analysed using a Fortessa flow cytometer (BD Biosciences). NKX2-1+ cells were isolated on the basis of NKX2-1-GFP expression, or expression of CD47 and CD26. For the latter, cells were stained with antibodies (CD47-PerCPCy5.5 #323110 and CD26-PE #323110, BioLegend) for 30 minutes on ice, and then CD47^hi^/CD26^low^ cells isolated by FACS ^23^. Where indicated, SFTPC+ iAT2s were purified by FACS on the basis of a SFTPC-tdTomato reporter gene ^24^ or on the basis of CPM expression (Novachem, 014-27501) ^29^. Cells were resuspended in sort buffer (Hank’s Balanced Salt Solution [ThermoFisher], 2% FBS, 10 mm Y-27632). Live cells were sorted using 10 mM calcein blue AM (Life Technologies), Zombie NIR or Zombie R718 (Biolegend 423106 and 423115). Cells were isolated using a FACS Aria (BD Biosciences) at the Murdoch Children’s Research Institute Flow Cytometry Core Facility.

To assess iMac differentiation or activation, cells were stained CD45-BV421 (Biolegend, 304032), CD14-PECy7 (Biolegend, 301814), HLA-DR-FITC (BD Bioscience, 347363), CD86-APC (Biolegend, 305412) and CD11b-APC (BD Bioscience, 550019) antibodies. iMac proliferation was measured by CFSE dilution using flow cytometry as per manufacturer’s instructions (ThermoFisher Scientific, C34554). Live cells (assessed by calcein blue AM or Zombie) were analysed on a Fortessa (BD Biosciences).

### Immunostaining and confocal imaging

Samples were fixed with 4% PFA (Santa Cruz, sc281692) for 20 minutes at room temperature and stored at 4°C prior to staining. ALI membranes were excised with a scalpel blade prior to immunostaining. Samples were permeabilised with 0.3% TritonX-100 (Sigma-Aldrich, T8787) and blocked with 4% normal donkey serum or normal goat serum (Sigma-Aldrich). Blocking solution was used to dilute primary antibodies which were incubated at 4°C overnight. Samples were washed, prior to incubation with fluorescently conjugated antibodies and counterstain with DAPI (Sigma-Aldrich, D9542) for 1 hour at room temperature. Live cell imaging or immunostaining were imaged with a LSM 900 confocal microscope (Zeiss) and images were processed using ImageJ. Antibodies used in this study were: SPC (Santa Cruz, sc518029), CD68 (Abcam, ab213363), HT1-56 (Terrace Biotech, TB-29AHT1-56), RSV (Abcam, ab43812 and Merck, AB1128). Fluorescently conjugated secondary antibodies were purchased from Invitrogen.

### Viral isolation and infection

Respiratory syncytial virus (RSV) strain A2 (ATCC VR-1540) was propagated in Hep-2 cells as previously described ^41^. Briefly, Hep-2 cells were infected at low multiplicity of infection (MOI) and incubated at 37°C for 5 days in DMEM (ThermoFisher, 21969035) supplemented with 10% foetal bovine serum (ThermoFisher, A5670701) and 10 U/mL penicillin, and 10 U/mL streptomycin (ThermoFisher, 15070063) (“DMEM complete”). Infected cells were scraped into the supernatant, then centrifuged at 1500 x g for 10 minutes. Clarified supernatant was underlaid with 5 ml sucrose cushions (30% (*m*/*v*) sucrose, 1X PBS, pH 7.4) in SW28 ultracentrifuge tubes and the virus pelleted at 20,000 x g for 90 minutes at 4°C. Pellets were resuspended in DMEM without supplementation and aliquots snap frozen on dry ice then stored at −80°C. Titres of RSV stocks or experimental samples were determined using an immunoplaque assay. In brief, Hep-2 cells were seeded in 96-well culture plates and inoculated with serial 1:5 dilutions of stock/sample for two hours with occasional agitation, inoculum removed then incubated for 3 days with a 1% methyl cellulose (Sigma-Aldrich, M7027) overlay in DMEM complete. Cells were fixed with 4% PFA for 15 minutes, permeabilised with 0.3% TritonX-100 for 15 minutes and washed three times in PBS-T (0.1% Tween 20). Fixed plates were blocked with 4% BSA for 1 hour, incubated with anti-RSV antibody (1:500) (Merck, AB1128) for 90 minutes, washed as before, then incubated with Alexa Fluor488 conjugated secondary antibody (ThermoFisher, A32790) for 1 hour. All steps were performed at room temperature. Plaques were manually counted on a fluorescent microscope (Zeiss Observer.Z1) at 10X magnification and titres calculated as plaque forming units per ml.

Influenza A virus (A/ PR8/34; H1N1) stocks were propagated in the allantoic cavity of 10-day embryonated chicken eggs through collaboration with the WHO Collaborating Centre for Reference and Research on Influenza (WHO CCRRI) at The Doherty Institute, Melbourne, and titred using MDCK cells by standard plaque assay, as previously described ^42^.

ALIs were infected apically with RSV-A2 MOI 10, or 1 or IAV H1N1 MOI 2 diluted in 50 uL DMEM. iMacs alone were cultured in tissue-culture treated 96 well plates for infection studies. Inoculum was removed after 2 hours, cells washed, then returned to air (ALIs) or media replenished (iMacs alone). ALI Apical washes with PBS were taken every 2-3 days to harvest shed virus. TEER measurements were taken simultaneously. To inhibit VEGFR2/KDR, 10 uM semaxinib (SU5416) (MedChemExpress, HY-10374) was added to the basolateral compartment after the initial viral inoculum was removed, and media replenished after 48 hours.

### Quantitative real-time PCR

RNA was extracted using the ISOLATE II RNA Mini Kit (Bioline, BIO-52073), as per manufacturer’s protocol. Complementary DNA (cDNA) was reverse transcribed using the Tetro cDNA Synthesis Kit (Bioline, BIO-65043). Quantitative real-time PCR (qRT-PCR) was run for 45 cycles using PowerTrack SYBR Green Master Mix (ThermoFisher, A46111) and custom primers (Table 1). For each biological replicate, the average Ct value for technical triplicates was calculated and normalised to the housekeeping gene (*ACTB*). Fold change was determined using 2^ΔΔCt.

**Table 1.**
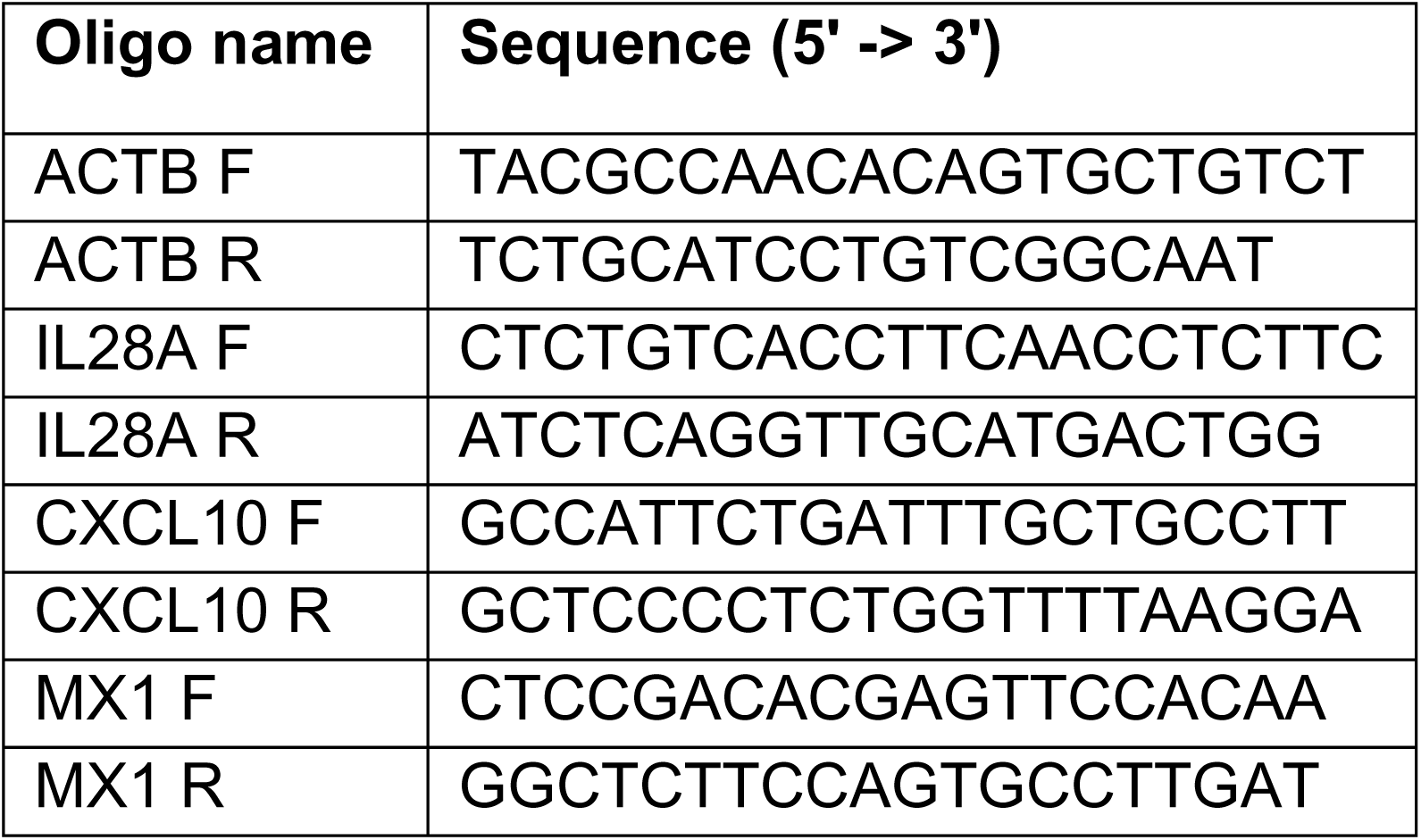
Oligonucleotide sequences used for qRT-PCR.

### Single-cell RNA-sequencing

Single-cell RNA-sequencing was performed using the Flex Gene Expression assay (10X Genomics, 1000475). iAT2 ALIs were established at air, and in some Transwells iMacs were added to the apical surface, as described above. 3 days later half the Transwells were infected with RSV (MOI 10), as described above. iMacs alone were plated on 6-well tissue-culture treated plates in CK-DCI, and infected as above. Uninfected or infected samples were dissociated 48 hours later using Accutase (Stem Cell Technologies, 7922). To achieve sufficient cell numbers Transwells from the same condition were pooled as appropriate. Cells were stained with Zombie R718 for 15 minutes at room temperature, then incubated with anti-CD45 antibody. Cells were then fixed with “Fixation buffer” (10X Genomics, 1000475) for 21 hours. After quenching fixation as per manufacturer’s instructions, Zombie-negative cells were sorted. In iAT2-iMac cocultures, CD45-iAT2s and CD45+ iMacs were sorted separately, then pooled after sorting 10:1. Cells were collected from the sort in Lo-bind Eppendorf tubes supplemented with RNAse inhibitor (Promega, M6101). Cells were counted then stored at −80°C, as per manufacturer’s recommendations.

After storage, monocultured iAT2s and iMacs, as well as infected iAT2s and iMacs were pooled 1:1 then all samples hybridised with Human WTA Probes BC001-BC004 as well as 26 custom spike-in probe pairs against RSV A2 genes at 40 nM per probe for all samples (Supplementary Table 1). Custom probes were designed according to 10X tech note CG000621_RevC and purchased as standard desalted oPools from IDT and resuspended in low EDTA TE buffer (ThermoFisher Scientific, #12090015). Samples were multiplexed prior to GEM generation using a Chromium iX (10X Genomics). GEMs were recovered and gene expression libraries constructed, following manufacturer’s protocols. Libraries were sequenced on an Illumina NovaSeq X plus (AGRF).

Sequencing generated reads with 94% >Q30. Single-cell RNA-sequencing bioinformatics analysis was performed using the Cellranger pipeline (version 8.0.0) was used to create fastq files and count matrices. Seurat (v5) was used for further analysis and data visualisation. Doublets and cells with more than 5% of reads mapping to mitochondrial genes were filtered out and data was normalised using SCTransform. Uniform Manifold Approximation and Projection (UMAP) and PCA were used for dimensionality reductions and clusters determined by the Louvain algorithm. Cell cycle stage was calculated, as described ^43^. Monocultured iAT2s and iMacs as well as infected monocultured iAT2s and iMacs combined prior to barcoding were assigned unique identities based on Louvain clustering and expression of NKX2.1 >1 and CD68 >1 respectively and analysed as unique biological samples thereafter. Differentially expressed genes were determined using the FindAllMarkers function implemented in Seurat, with a Wilcoxon Rank Sum test and a default log fold change of 0.1. Gene set enrichment analysis (GSEA) was performed with hypeR using ranked differentially expressed gene lists ^44^. Cell-type identification was performed using scType ^45^. Cell-cell communication was inferred using CellChat ^46^. Data are deposited on GEO.

Analysis of previously published datasets was performed to analyse expression of *CSF1* and *CSF2* in primary AT2s and iAT2s. Healthy AT2s were subsetted based on expression of *SFTPC* (>4) from a previously published study of the human lung ^47^. Control iAT2s from previous studies ^30,31^ were subsetted for further analysis here.

### Lipid uptake assay

To label intracellular lipids, iAT2s were plated on Matrigel (Corning #354277)-coated plates then incubated with 5 µg/mL FM™ 4-64 Dye (N-(3-Triethylammoniumpropyl)-4-(6-(4-(Diethylamino) Phenyl) Hexatrienyl) Pyridinium Dibromide) (ThermoFisher Scientific, T13320) for 20 minutes. iAT2s were washed prior to addition of a ‘secretagogue cocktail’ consisting of 100 nM ATP (ThermoFisher Scientific, R0441) and 300 nM PMA (Abcam, AB147465). iMacs were immediately added. At 20 or 90 minutes later, cultures were dissociated with Accutase, stained for cell surface markers on ice and analysed on a Fortessa flow cytometer (as above).

### Wound healing assay

300,000 iAT2s were plated on Matrigel (Corning #354277)-coated 48-well plates for 24 hours, prior to the addition of iMacs (10:1 ratio). A scratch through the iAT2 monolayer was introduced (time 0) and imaged over 48 hours, as we have previously described ^31^. The area of the scratch was calculated using ImageJ and calculated relative to the area at time 0.

### Statistics

Statistical analyses were performed using unpaired two-tailed Student’s t tests for comparisons between two groups and one-way analysis of variance (ANOVA) with a Tukey multiple comparisons test for comparisons among three or more groups. Details for each analysis are provided in the figure legends. Data are presented as means, with error bars representing standard deviation (SD). A P value of <0.05 was considered statistically significant, with annotations on graphs as follows: *P < 0.05, **P < 0.01, ***P < 0.001, ****P < 0.0001.

## Results

### Establishment of iPSC-derived AT2s and macrophages coculture at air-liquid interface

To develop an advanced model of the human alveolus, we constructed a coculture system that combined iPSC-derived AT2s and macrophages in the context of physiologically relevant ALI culture system. Since each cell type arises from distinct germ layers and regions of the embryo, there are no established differentiation protocols to concurrently derive both cell types. Thus, to ensure our model contained pure, well-characterised cell populations, we separately differentiated iAT2s ^22,24^ and iMacs ^25^, using previously published defined conditions (Figure 1A and Supplemental Figure 1A). As expected, prior to coculture, iAT2s expressed NKX2-1 and surfactant protein C (SPC) ^23,24^, while iMacs expressed CD14, CD11b and CD68 ^25^ (Figure 1B). To establish cocultures, iAT2s were matured in an ALI culture, as we have previously described ^22,26,29^, prior to the addition of macrophages in the apical compartment (Figure 1A). Cocultures were maintained in iAT2-media (“CK-DCI”). Live cell imaging of co-cultured iAT2s (based on endogenous SFTPC-tdTomato ^24^) and iMacs (labelled with CellTrace dye) demonstrated dynamic movement, viability and longevity of iMacs in the cocultures (Figure 1C, Supplemental Figure 1B and Video 1). Confocal imaging of stained cocultures indicated a close association between iAT2 and iMacs, with macrophages localised to the apical surface only (Figure 1D). We found that a similar approach could be employed to coculture iAT2s and primary alveolar macrophages from cryopreserved bronchoalveolar lavage fluid at air-liquid interface (Supplemental Figure 1C). To investigate whether an analogous coculture could be established with AT1 cells, we created iAT1 cultures at ALI ^36^. Compared with iAT2-iMac cocultures, we observed very few adhered iMacs to the iAT1s (Supplemental Figure 1D).

**Figure 1.**
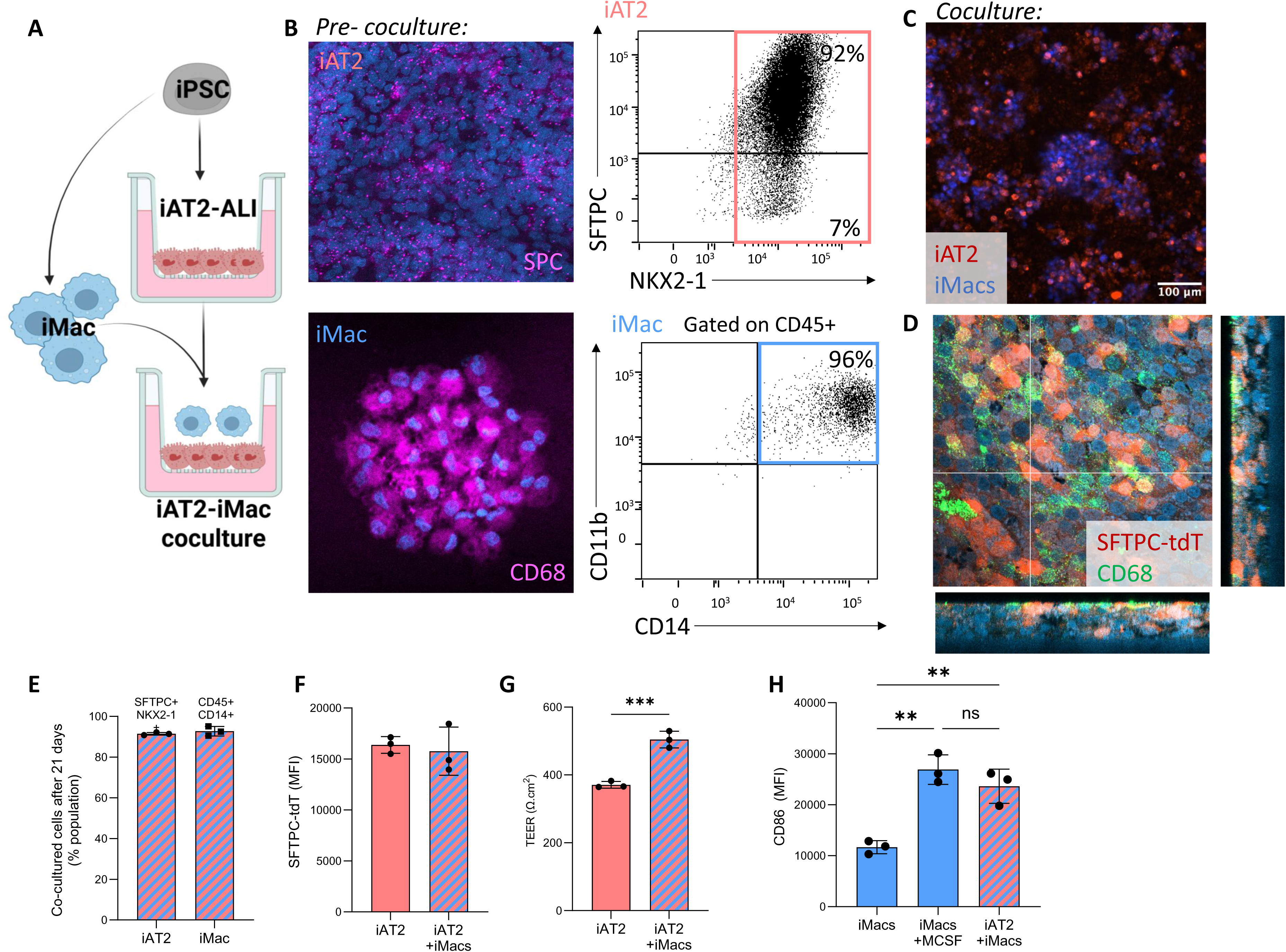
Establishment of iPSC-derived AT2 and macrophage air liquid interface cocultures. A) Schematic representation of separate iAT2 and iMac differentiations, iAT2s plated at air-liquid interface (ALI) for 3-7 days prior to addition of iMacs to the apical compartment. B) Prior to coculture iAT2 express surfactant protein C (SPC) protein (top, left), SFTPC-tdTomato and NKX2-1*-*GFP (top, right). iMacs express CD68 (bottom, left), CD14 and CD11b (bottom, right). C) Live cell confocal imaging of iAT2 (red, marked by SFTPC-tdTomato) and iMacs (blue, stained with CellTrace Violet) in coculture at ALI. D) Confocal imaging of iAT2s (red, SFTPC-tdTomato) and iMacs (green, CD68). E) Summary of flow cytometry indicating that following 21 days coculture showing that iAT2s maintain expression of SFTPC-tdTomato and NKX2-1-GFP and whilst iMacs express CD45 and CD14. F) Mean fluorescence intensity (MFI) of SFTPC-tdTomato in iAT2 alone, or iAT2s after coculture with iMacs 7 days of coculture. G) Transepithelial electrical resistance (TEER) in iAT2 alone, or iAT2s cocultured with iMacs for 7 days. H) MFI of CD86 after 7 days cultured in CK-DCI, CK-DCI + M-CSF, or with iAT2s in CK-DCI. n = 3 experimental replicates of independent wells of a differentiation; error bars represent SD. Statistical significance was determined by unpaired, two-tailed Student’s t test (two groups) or one-way ANOVA (>2 groups); *p < 0.05, **p < 0.005, ***p < 0.001.

To assess the effect of coculture on the identity of either cell type, cocultures were dissociated three weeks post establishment and cells analysed by flow cytometry. Both iAT2s (SFTPC+ NKX2-1+) and iMacs (CD45+ CD14+) retained their original identity (Figure 1E), although at this stage, iMacs comprised 1% of total cell numbers, compared to the initial seeding proportion of 10% (Supplemental Figure 1E). iAT2 expression of SFTPC-tdTomato was not significantly altered between cells in iAT2 mono- vs cocultures (Figure 1F). Interestingly, barrier integrity, measured by transepithelial electrical resistance (TEER), was significantly enhanced by coculture with iMacs (Figure 1G). Levels of the costimulatory receptor, CD86, was upregulated on cocultured iMacs (Figure 1H and Supplemental Figure 1F-G), which was comparable to that observed with iMacs derived from cultures supplemented with macrophage colony stimulating factor (M-CSF) (Figure 1I). Collectively, these data demonstrate the successful establishment of iAT2–iMac cocultures at ALI, preserving cell identity while enhancing the functional properties of both cell types.

### Coculture establishment is unaltered by route of iMac directed differentiation

Although the alveolar macrophages in the lung at birth arise from primitive haematopoiesis, these are gradually replaced by monocyte-derived macrophages which result from definitive haematopoiesis ^48,49^. Numerous protocols describing the generation of iMacs have been established ^50^, with primitive vs definitive-derived iMacs displaying some functional differences *in vitro* ^51^. Since the iMacs we had used thus far arose from a protocol that excludes retinoids and is therefore likely to generate macrophages mirroring those derived from extraembryonic (primitive) haematopoiesis ^25^, we also sought to examine the characteristics of macrophages generated with a protocol that supported the development of intra-embryonic (definitive) haematopoietic stem/progenitor cell intermediates ^38,52^. iMacs derived through primitive and definitive routes both responded to respiratory syncytial virus (RSV) infection, upregulating interferon stimulated genes (*IFIT2* and *MX1*), although definitive iMacs displayed a higher magnitude of response late in infection (Supplemental Figure 2A-C). We next created iAT2-iMac co-cultures using iMacs derived from definitive haematopoiesis. As observed with our cocultures containing primitive iMacs, cocultures formed with definitive iMacs maintained their identity for 7 days (Supplemental Figure 2D-E). Taken together, these findings suggest that the route of iMac differentiation does not significantly impact the establishment of cocultures.

### iPSC-derived macrophages promote surfactant gene expression by iAT2s

Surfactant homeostasis in the lung is maintained through synthesis and recycling by AT2s and degradation by alveolar macrophages ^53^. To explore how coculture with iMacs would influence the transcriptome of iAT2s, we performed single-cell RNA-sequencing (scRNA-seq) of monocultures and cocultures (Figure 2A and Supplemental Figure 3A-D). iAT2s were established in air for two days, prior to the addition of iMacs for five days. Uniform manifold projection (UMAP) visualisation suggested large overlap of transcriptomes of iAT2s alone and cocultured iAT2s (Figure 2A), although unsupervised cell clustering (Louvain) revealed a cluster was predominated by cocultured iAT2s (cluster 2) (Figure 2B-C). Importantly, coculturing iAT2s did not adversely alter expression of key iAT2 markers, nor promote the emergence of non-lung endoderm, off-target lineages (Supplemental Figure 3C). Indeed, differential gene expression analysis revealed an upregulation in AT2 genes (*PGC, SFTPB, ABCA3*) and proliferation genes (*MIKI67, TOP2A*) in cocultured iAT2s, compared to iAT2s alone (Figure 2D). Gene set enrichment analysis revealed pathways involved in lipid synthesis and metabolism, glycosylation, and ABC transporters were upregulated in iAT2s following coculture with iMacs (Figure 2E and Supplemental Figure 3E-F), suggestive of changes in surfactant synthesis and processing. N-linked glycosylation is a key posttranslational modification, essential for the secretion and function of surfactant proteins ^54,55^. Intriguingly, expression of the surfactant proteins that undergo glycosylation, *SFTPB, SFTPA1,* and *SFTPD,* were upregulated in cocultured iAT2s (Figure 2F). Furthermore, *ABCA3*, the essential surfactant lipid transporter on lamellar bodies, was amongst the differentially upregulated genes in iAT2s after iMac coculture (Figure 2G). Quantification of AT2 differentiation and maturation gene modules ^56^ demonstrated iAT2 maturation was significantly increased in iAT2s in cocultures, compared with iAT2s alone (Figure 2H and Supplemental Figure 3G). Collectively, our single-cell transcriptomic profiling suggested that iAT2s increase surfactant production and packaging in the presence of iMacs, potentially as a consequence of surfactant metabolism by the macrophages. To confirm that iMacs were taking up surfactant in our model, we labelled lipids with a lipophilic dye and then stimulated iAT2s to secrete surfactant, as previously described ^28^. Flow cytometry revealed that iMacs rapidly and efficiently phagocytosed extracellular lipids secreted by iAT2s (Figure 2I and Supplemental Figure 3H). Importantly, this was not due to cell death of iAT2s (Supplemental Figure 3I), suggesting that iMacs were metabolising surfactant in our iAT2-iMac cocultures system, mirroring the activity of their *in vivo* counterparts.

**Figure 2.**
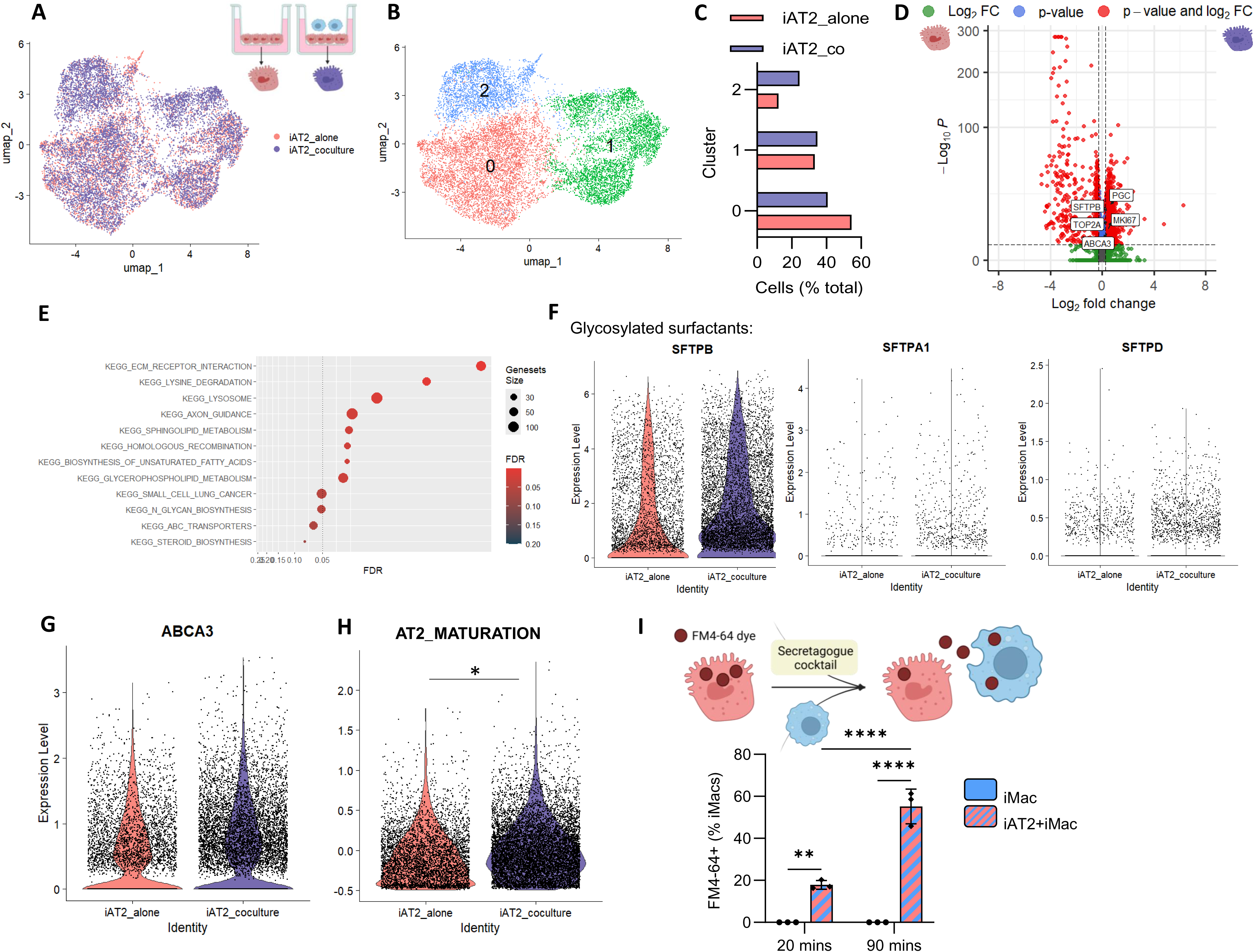
Coculture with iMacs promotes iAT2 transcriptional signature. A) Uniform manifold projection (UMAP) of iAT2 cells alone (pink) and iAT2 cells following coculture with iMacs (purple). B) Louvain clustering at a resolution of 0.1. C) Proportion of iAT2 alone (pink) or iAT2 after coculture (purple) in each cluster at Louvain resolution 0.1. D) Volcano plot of differentially expressed genes upregulated in iAT2 alone (pink, left) or iAT2 after coculture (purple, right). E) Upregulated pathways from gene set enrichment analysis in iAT2s after coculture. F) Violin plots of differentially expressed genes included *SFTPB, SFTPA1*, *SFTPD* and G) *ABCA3*. H) Module score of AT2 maturation gene set ^56^ indicating that culture enhances AT2 maturation. I) iAT2s were treated with the lipophilic dye, FM4-64, washed then treated with a secretagogue cocktail (ATP and PMA). iMacs were immediately added and incubated for 20 or 90 minutes, prior to collection and flow cytometry to measure internalised FM4-64 in iMacs. n = 3 experimental replicates of independent wells of a differentiation; error bars represent SD. Statistical significance was determined by a Welch Two Sample t-test or a two-way ANOVA; *p < 0.05, **p < 0.005, ***p < 0.001.

### iPSC-derived macrophages are maintained through iAT2-derived M-CSF

We next sought to examine the consequences of coculture on iMacs. scRNA-seq revealed that cocultured iMacs cluster discretely from mono-cultured iMacs (Figure 3A-B and Supplemental Figure 4A). Coculture did not cause iMacs to enter cell cycle (Figure 3C). Yet, coculture of iMacs did significantly alter their transcriptome, compared to iMacs alone (Figure 3D). To compare cocultured iMacs to *in vivo* cell types, we used scType for cell-type identification ^45^. Cocultured iMacs were most similar to alveolar macrophages, compared with iMacs cultured alone, which were classified broadly as immune system cells in the lung (Figure 3E). In light of the “tissue-resident”-like identity that cocultured iMacs adopted, we next sought to understand the influence of iAT2s in this process. In the adult human lung, AT2s produce M-CSF and granulocyte-macrophage colony stimulating factor (GM-CSF) ^57^, and we observed that iAT2s express similar levels of both cytokines (Figure 3F). Strikingly, in the presence of iMacs, iAT2s further upregulated expression of M-CSF (encoded by *CSF1*) and to a lesser extent GM-CSF (encoded by *CSF2*) (Figure 3G and Supplemental Figure 4B). We next assessed cell-cell interactions using CellChat ^46^ between cocultured iAT2s and iMacs. Among the significantly enriched signals from iAT2s to iMacs was CSF1-CSF1R (Figure 3H and Supplemental Figure 4C), suggesting that iMacs may be sustained in coculture by iAT2-derived M-CSF. Given M-CSF is soluble we reasoned that iMac maintenance would be independent of contact with iAT2s. To test this, we collected conditioned media from iAT2 cultures and used this to supplement isolated iMac cultures. Conditioned media did not alter iMac proliferation but significantly upregulated HLA-DR expression, to a similar extent as iMacs maintained in M-CSF containing media (Figure 3I and Supplemental Figure 4D). Of note, GM-CSF supplemented iMacs expressed significantly higher levels of HLA-DR, consistent with enhanced activation mediated by GM-CSF in previous reports ^58,59^. To understand whether iAT2-derived M-CSF was directly responsible for iMac activation, we used neutralising antibodies to block M-CSF, GM-CSF or both in iAT2-conditioned media. Blockade of M-CSF or both M-CSF/GM-CSF, but not GM-CSF alone, inhibited the upregulation in HLA-DR expression in iMacs (Figure 3J). Collectively, our data suggest that iMacs are maintained in cocultures by iAT2-derived M-CSF.

**Figure 3.**
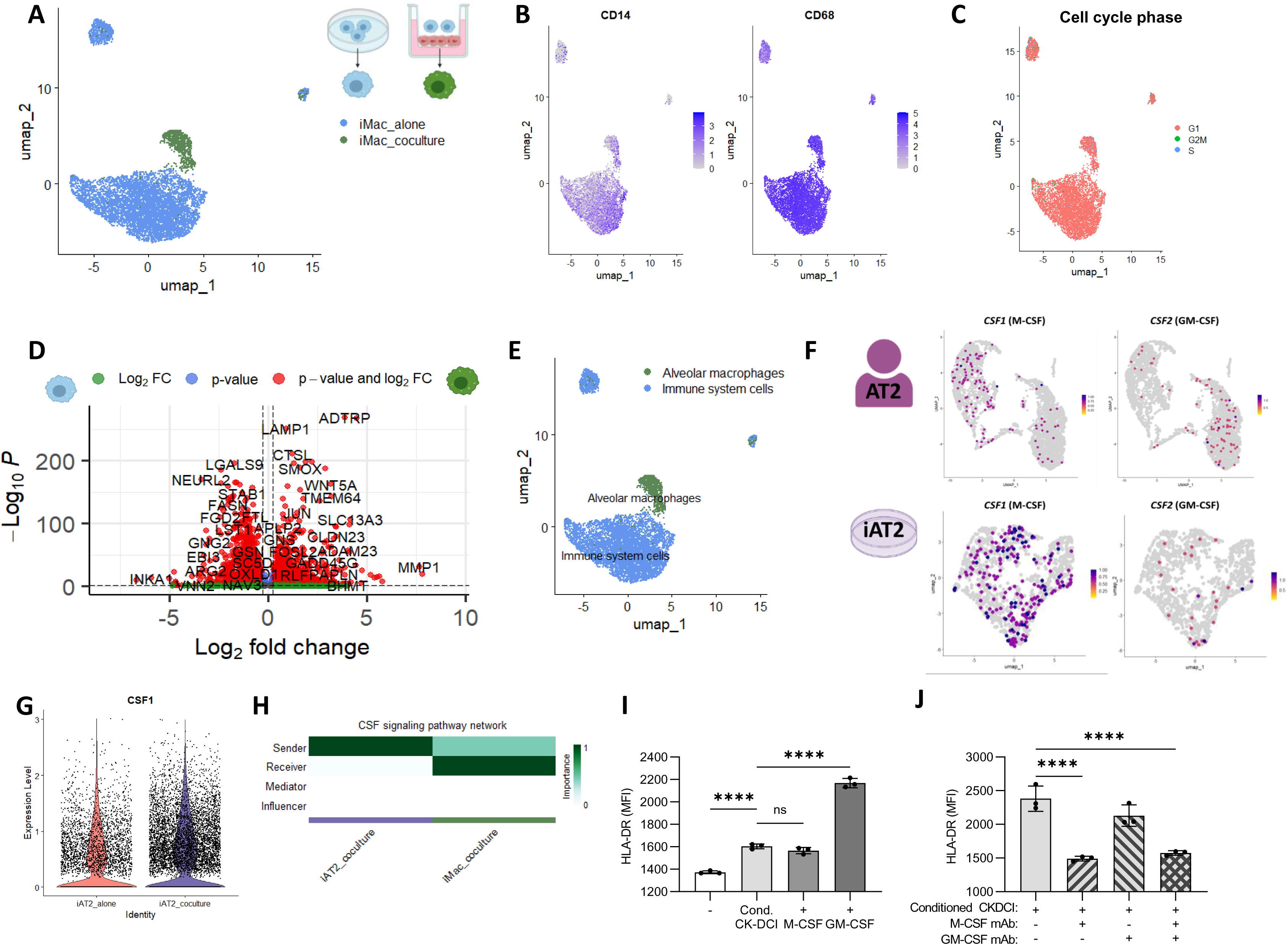
iMacs are sustained in coculture by iAT2-derived M-CSF. A) Uniform manifold projection (UMAP) iMacs alone (blue), or iMacs following coculture with iAT2s (green) showing distinct clustering based on original identity B) UMAP showing expression of *CD14* and *CD68*. C) UMAP of cell cycle phase. D) Volcano plot of differentially expressed genes upregulated in iMac alone (blue, left) or iMac after coculture (green, right). E) scType analysis of iMacs alone, or iMacs following coculture with iAT2s. F) *CSF1* and *CSF2* expression in adult human AT2s ^47^ and iAT2s ^30,31^. G) *CSF1* expression in iAT2 alone (pink), or iAT2 following coculture with iMacs (purple) showing upregulation of *CSF1* expression by iAT2s in coculture. H) CSF signalling pathway identified in CellChat analysis of iAT2s in coculture (purple) or iMacs in coculture (green). I) Conditioned CK-DCI from iAT2s was added to iMacs alone. Alternatively, iMacs were cultured in CK-DCI supplemented with M-CSF or GM-CSF. Mean fluorescent intensity (MFI) of HLA-DR was measured by flow cytometry after 72 hours. J) iMacs treated with conditioned CK-DCI from iAT2s were incubated with neutralising antibodies against M-CSF, GM-CSF or both. HLA-DR MFI was measured by flow cytometry after 72 hours.

### iPSC-derived macrophages promote inflammation and antiviral immunity following respiratory viral infection of iPSC-derived type 2 alveolar epithelium

While most respiratory viruses preferentially replicate in the upper respiratory tract, these can spread to the lower respiratory tract and alveolus, causing severe infections like pneumonia. In the alveolus, human-tropic respiratory viruses like RSV and IAV commonly target AT2s ^10,11^. To test our coculture platform in the context of infection, we treated cultures with the viral mimetic, poly(I:C), or infected with IAV or RSV. Following exposure, iAT2-iMac cocultures expressed significantly higher antiviral interferon (IFN)-λ and antiviral interferon-stimulated genes (ISGs) (*CXCL10* and *MX1*), compared to iAT2s alone or iMacs alone (Supplemental Figure 5A-D). To explore this further, we performed scRNA-seq 48 hours after RSV infection of iAT2s or iMacs alone, or iAT2-iMac cocultures (Figure 4A and Supplemental Figure 5E-G). UMAP visualisation and cell clustering revealed that iMacs segregated from iAT2s following infection. Moreover, cocultured iMacs clustered distinctly from iMacs alone (Figure 4A-B). Although this division was less defined in the iAT2s, we did observe certain clusters (2 and 7) predominantly contained iAT2s from cocultures (Figure 4B). Three clusters (8, 9 and 12) contained RSV transcripts, representing actively infected iMacs alone, infected iMacs from cocultures and infected iAT2s, respectively (Figure 4C). Previous studies in mice have shown alveolar macrophages are key producers of type I interferons (IFN) ^60^, the proinflammatory cytokines tumour necrosis factor (TNF), interleukin (IL)-1α, IL-1β, and IL-6, and chemokines (e.g., C-C motif ligand [CCL]-2-4) ^61,62^ during RSV infections. Supporting the fidelity of our coculture system, iMacs were the primary producer of these antiviral cytokines, proinflammatory cytokines, and chemokines (Supplemental Figure 5H-I). Furthermore, unbiased gene set enrichment analysis revealed that inflammatory pathways were enriched in iMacs, compared to iAT2s (Figure 4D-E). Overall, while these pathways were upregulated in both iMacs alone and in coculture, cocultured iMacs appeared less inflammatory yet enriched for more viral transcripts (Figure 4F), suggesting the coculture environment shapes iMac immune responses.

**Figure 4.**
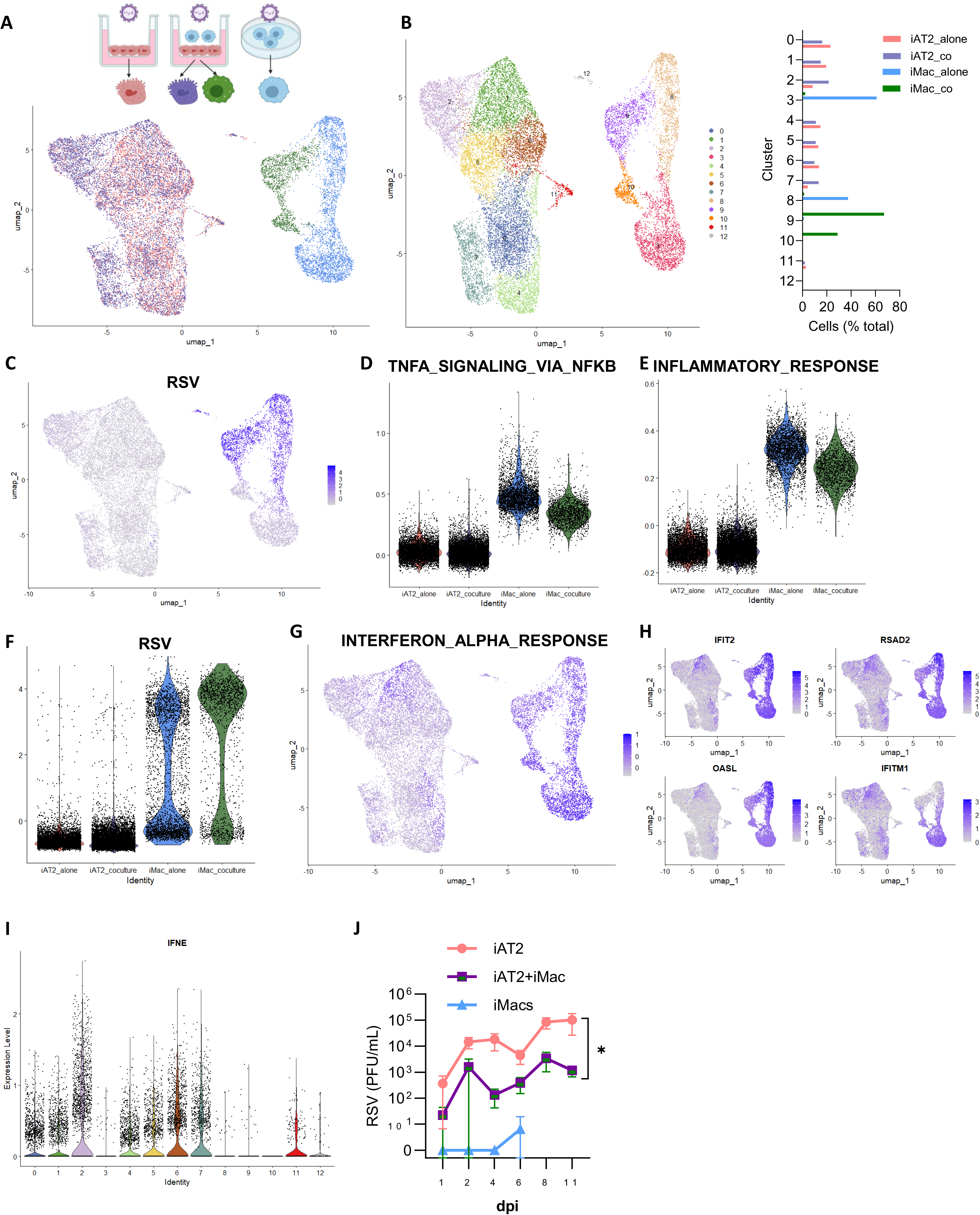
iMacs augment inflammation and antiviral immunity during viral infections of cocultures. A) Single cell RNA-seq analysis of iAT2 alone, iMac alone and cocultures 48 hours following RSV (MOI 10) infection. Uniform manifold projection (UMAP) of cells showing original identity. B) Louvain clustering at a resolution of 0.4. C) RSV transcripts. D-E) Gene sets enriched in iMacs following coculture from the Hallmark database. F) RSV transcripts (whole genome). G) Interferon Alpha Response from the Hallmark database. H) Interferon stimulated genes. I) IFNE expression in clusters (resolution 0.4, panel B). J) Shed RSV in the apical compartment of ALIs containing iAT2 alone or iAT2-iMac cocultures was measured by plaque assay.

Innate antiviral immunity is a vital defence against respiratory viruses through the production of type I and III IFNs which induce ISGs that directly inhibit viral replication or recruit immune cells. Innate IFN responses, including ISGs, were more enriched in iMacs compared with iAT2s following infection (Figure 4G-H). Interestingly, expression of certain ISGs (e.g., IFITM1) were largely restricted to bystander iMacs and iAT2s (i.e., cells that did not express RSV transcripts in virus infected cultures). Although actively infected iAT2s were rare, we sought to understand whether coculture with a professional immune cell would reshape antiviral immunity in iAT2s. Notably, certain clusters of bystander iAT2s, but not the actively infected iAT2s, were enriched for IFN signalling (Figure 4G-H). Cluster 2, which predominantly contained iAT2s from cocultures, was significantly enriched for type I IFN (*IFNE*) as well as IFN receptors (*IFNLR1*) and downstream signalling molecules (*TYK2*) (Figure 4I and Supplemental Figure 4J). This indicates that in iAT2-iMac cocultures, both cell types have augmented antiviral immunity, suggesting that viral replication and shedding may be partially constrained in this platform. To investigate this, we infected iAT2 or iAT2-iMac ALIs, or submerged iMacs alone with RSV, and measured shed live virus over 11 days. RSV release was almost entirely absent in infected iMacs alone, consistent with previous reports showing abortive infection of RSV in alveolar macrophages ^63^, although these cells died after six days (Figure 4J). Virus continued to shed in iAT2 cultures over 11 days, and this was significantly reduced in cocultures with iMacs (Figure 4J), consistent with enhanced antiviral immunity restraining RSV replication.

### iPSC-derived AT2 repair is influenced by macrophages

Alveolar macrophages play critical roles in wound healing after lung injury, by stimulating proliferation of structural cells (epithelium and fibroblasts), promoting angiogenesis, and dampening inflammation ^64^. To explore whether iMacs would influence wound healing, we performed a scratch assay through a monolayer of iAT2s ^31^ with or without iMacs, then monitored wound closure. iMacs significantly accelerated iAT2 wound closure over a 48-hour period (Figure 5A-B). Since respiratory viruses can injure the epithelium through disrupted tight junctions and cytotoxicity ^65,66^, we also assessed whether iMacs would change transepithelial electrical resistance (TEER) during RSV infection of iAT2s. By 11 days post infection, iAT2-iMac cocultures display improved barrier integrity, compared to iAT2s alone (Figure 5C). Interestingly, early in infection (2 days post infection) the presence of iMacs impaired TEER (Figure 5C). To investigate pro- or anti-repair mechanisms engaged by epithelial-macrophage crosstalk, we assessed cell-cell communication at 2 days post RSV infection in our scRNA-seq dataset. Infection prompted six new ligand-receptor interactions, which were not present between iAT2s and iMacs in uninfected conditions (Supplemental Figure 6A-C). Among these infection-only pathways, the VEGFA-VEGFR2 axis was the only one in which the damaged iAT2s acted as senders, expressing *VEGFA*, while iMacs served as direct receivers via VEGFR2 (also known as *KDR*), forming a linear ligand-receptor interaction (Figure 5D-G and Supplemental Figure 6D). Strikingly, iMacs further upregulated *KDR* expression during coculture, compared to iMacs alone (Figure 5G). VEGF signalling plays varied roles during infection and in recovery from lung injury, acting as a chemotactic agent, mitogen, and angiogenic factor ^67–71^. Whether VEGF has predominantly beneficial or detrimental effects, particularly in the context of human AT2 cells and macrophages during respiratory viral infections, is unclear. To investigate this, we inhibited VEGFR2/KDR signalling using semaxinib (SU5416) ^72^. RSV infection impaired TEER in iAT2s alone and this was unaltered by semaxinib treatment (Figure 5H). As we had observed previously (Figure 5C), TEER significantly declined in iAT2 cultures containing iMacs. However, by 4 days post infection, barrier integrity was entirely restored by inhibiting VEGFR2/KDR signalling in cocultures (Figure 5H). Unsurprisingly, viral shedding was not affected by semaxinib treatment (Supplemental Figure 6E). Together, this suggests that early during infection, VEGFA-VEGFR2 signalling between iAT2s and iMacs alters epithelial permeability.

**Figure 5.**
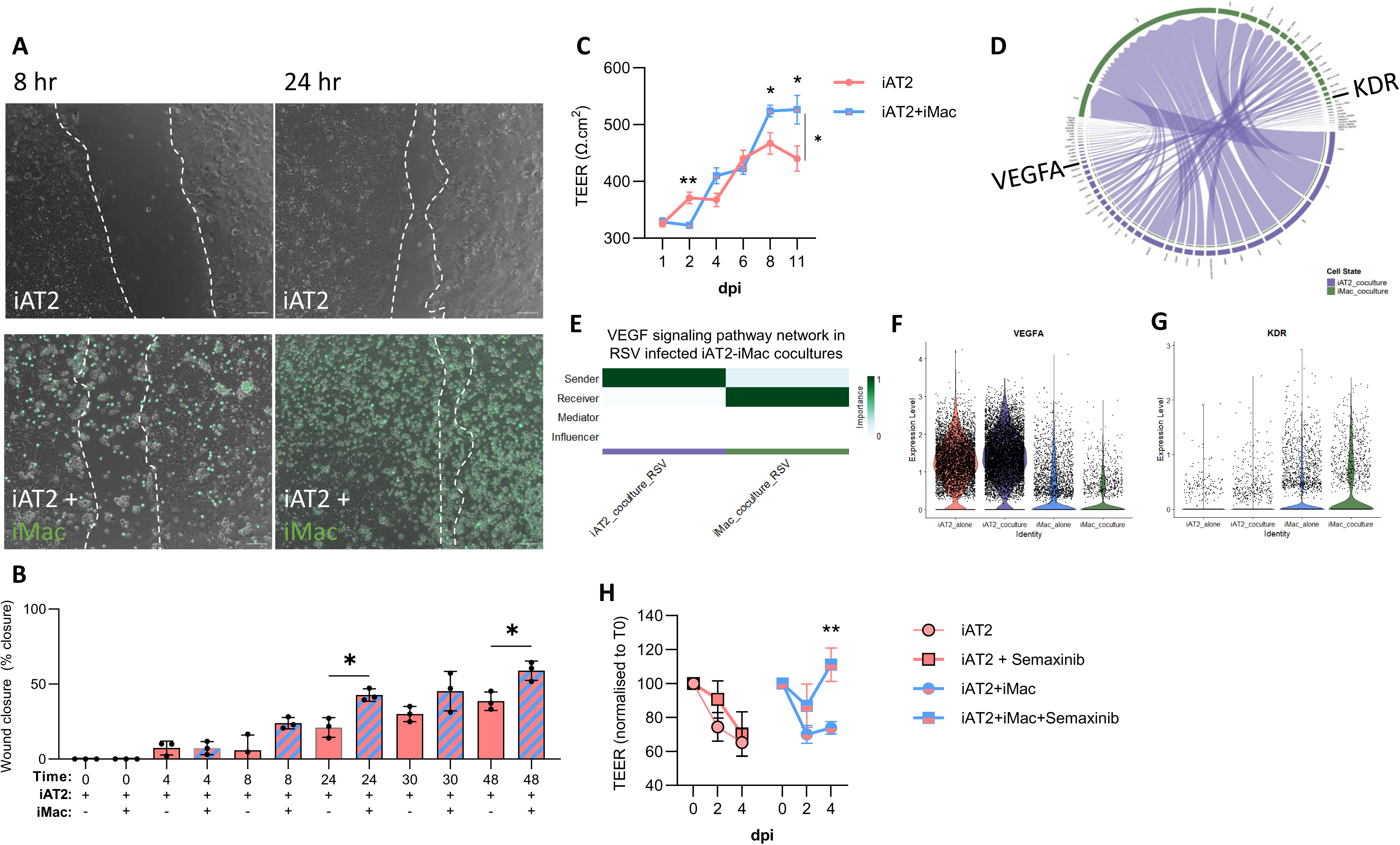
iMacs influence iAT2 repair through VEGF signalling. A-B) iAT2s were plated in 2D and allowed to reach confluence before a scratch wound was made. iMacs were labelled with CFSE (green) and added immediately after the scratch wound. Wound closure was calculated as a percentage of the initial wound over a 48-hour period. Scale bar = 100 um. C) iAT2s alone or iAT2-iMac cocultures were infected with RSV (MOI 10) and transepithelial electrical resistance (TEER) was measured over 11 days. D) Circos plot showing CellChat analysis of iAT2 signals (purple) to iMacs (green) in RSV infected cocultures. E) VEGF signalling pathway identified in CellChat analysis of iAT2s in coculture (purple) or iMacs in coculture (green). F) VEGFA and G) KDR (VEGFR2) expression in RSV-infected iAT2s alone, iAT2s from coculture, iMacs alone, or iMacs from coculture, showing iAT2s are the predominant source of VEGFA during RSV infection, and cocultured iMacs upregulate KDR. H) Transepithelial electrical resistance (TEER) of ALIs containing iAT2s alone or iAT2-iMac cocultures infected with RSV (MOI 10) then treated with semaxinib (KDR inhibitor) in the basolateral compartment. n = 3 experimental replicates of independent wells of a differentiation; error bars represent SD. Statistical significance was determined by a one-way or two-way ANOVA; *p < 0.05, **p < 0.005, ***p < 0.001.

## Discussion

Many acute and chronic respiratory diseases directly impact the alveoli. The development of human model systems that recapitulate interactions between key cell types will be critical for the discovery of new therapeutics. In this study, we establish the first iPSC-derived platform that incorporates AT2 cells and macrophages in a physiologically relevant air-liquid interface culture system which is easily amenable to infection studies. Coculture upregulated AT2-specific genes and lipid synthesis in iAT2s and iMacs phagocytosed surfactant. iAT2s supported iMacs in coculture through the production of M-CSF and iMacs adopted an alveolar macrophage-like phenotype. Tellingly, iMacs promoted proinflammatory signalling, antiviral immunity and limited viral replication during respiratory viral infections. Additionally, we found that iMacs influenced epithelial barrier repair and integrity, in part through VEGFA-VEGFR2 signalling.

Recent studies have described the incorporation or codevelopment of macrophages in iPSC-derived organotypic models, including the gut and brain ^73,74^. In these models, iMacs acquire transcriptional signatures resembling tissue-resident macrophages, regulate immune signalling, and promote organoid maturation ^73,74^. Mirroring these observations, iMacs in our model adopted an alveolar macrophage transcriptional signature, enhanced proinflammatory signalling and antiviral immunity during viral infections, and supported iAT2 maturation - features not observed in previous iPSC-derived alveolar models ^33,34^.

When establishing our model, we found that exogenous macrophage-supportive factors were unnecessary since iAT2s alone could sustain iMacs. During lung development the alveolar epithelium arises concurrently with alveolar macrophage differentiation ^49,75^, and alveolar epithelial cells remain a major source of GM-CSF and M-CSF into adulthood ^57^. iAT2s express both M-CSF and GM-CSF transcripts at levels comparable to *in vivo* human AT2s, with M-CSF expression further upregulated in cocultures. Of note, other macrophage-supportive cytokines like IL-3 and IL-34 were not expressed by iAT2s. iAT2-derived M-CSF appeared crucial to the maintenance of iMacs. We previously demonstrated that M-CSF alone is sufficient to induce and sustain functional iMacs ^25^, consistent with the ability of iAT2-derived M-CSF to maintain a stable population of iMacs within co-cultures. It is important to note that GM-CSF is indispensable for the development and survival of alveolar macrophages in mice ^49,57,76^, whereas M-CSF-deficient mice exhibit reduced alveolar macrophage numbers, which can be compensated by other cytokines ^77^. Thus, while M-CSF signalling appeared critical for the iAT2-iMac cocultures, we cannot entirely exclude a role for iAT2-derived GM-CSF.

Respiratory viruses commonly cause pneumonia in infants, with RSV a leading cause of pneumonia cases, hospitalisations and mortality in this age group ^78^. RSV is a human restricted pathogen, with minimal replication evident in small animal models ^79^. Furthermore, since respiratory viruses enter through the apical side of the epithelium, ALI models (but not organoids) support robust infection ^80^. To our knowledge, this is the first study to describe RSV infection in an ALI model that recapitulates the biology of primary AT2s, using both iAT2s alone and iAT2s with iMacs. Previous AT2 studies have primarily focused on other human-specific viruses such as coronaviruses ^26,81,82^. Surprisingly, despite productive and sustained infection, our scRNA-seq data revealed very few iAT2s were actively infected at 48 hours. Furthermore, only subsets of iAT2 bystanders activated antiviral defences, such as IFNs and ISGs, in contrast to RSV-infected airway epithelial cells, where antiviral responses appear more uniform ^83^. It would be interesting in future studies to assess whether a gradient of iAT2 bystander antiviral responses to RSV is determined by proximity to an infected cell ^84^.

During viral infections in mice, alveolar macrophage depletion can either lead to respiratory failure ^6^ or improve survival ^7,8^, suggesting potentially virus-specific effects and underscoring the delicate balance between inflammation, antiviral immunity, cell death, and repair. In our cocultures, iMacs were the primary infected cell type, aligning with findings from bronchoalveolar lavage studies of RSV-infected infants ^85,86^. Moreover, the presence of iMacs in cocultures significantly reduced viral replication. iMacs likely impaired viral burden through both abortive infection ^63,87^ and through prompting augmented antiviral immunity in iAT2s. This reiterates the importance of incorporating innate immune cells into respiratory epithelial models to faithfully recapitulate the sequelae of infection.

Alveolar epithelial damage during viral infections plays a key role in pneumonia pathology and can lead to severe complications such as acute respiratory distress syndrome ^13,14^. We demonstrated that iMacs promote iAT2 repair in both a scratch assay and during RSV infection. In mice, several macrophage-derived ligands, e.g., Wnt and IL-1β, have been implicated in AT2 proliferation, differentiation and recovery from injury ^88,89^. Our scRNA-seq analysis identified Wnt signalling between iMacs and iAT2s at baseline and during infection; however, we were unable to determine any potential role that this axis may play in repair due to the presence of a GSK3β inhibitor in the media, which is necessary for iAT2 maintenance ^24,90^. Interestingly, VEGF-A/VEGFR2 signalling emerged between iAT2s and iMacs only following viral infection, increasing barrier permeability early in infection. While macrophages are typically a source of VEGF in other organs, AT1s and to a lesser extent AT2s, are the main producers in the lungs ^21^, explaining *VEGFA* expression patterns in our cocultures. VEGF-C/VEGFR3 signalling is known to regulate macrophage functions such as efferocytosis during acute lung injury ^91^; however, this appears to be the first report of VEGF-A from human AT2s signalling through VEGFR2 on macrophages during a respiratory viral infection. Future studies should explore whether clinically used VEGF-targeting therapies, such as small molecules and monoclonal antibodies employed in cancer treatment, influence respiratory viral infection outcomes.

The development of iAT2-iMac ALI cocultures is an important first step towards more closely aligning the complexity of *in vitro* alveoli models with their *in vivo* counterparts. Ideally, these models will extend to incorporating other cell types, including AT1 cells. However, despite recent progress in generating pure iAT1 cells ^36^, current methods are unable to generate populations of iAT2s and iAT1s at frequencies that yield the proportion of each cell type present within native alveolus. Similarly, our model could be further elaborated to include other lung resident immune cells, such as dendritic cells, innate lymphoid cells, and tissue-resident lymphocytes. However, as a starting point, our iPSC-derived platform serves as a human *in vitro* model for studying AT2-macrophage interactions in homeostasis, infection with human-tropic respiratory viruses, and repair. Access to this model should facilitate disease modelling by providing insights into the relative contributions of AT2s and alveolar macrophages to respiratory disease initiation and progression.

## Acknowledgements

We are grateful to members of the Lung Disease, Immune Development, and Blood Development laboratories at the Murdoch Children’s Research Institute (MCRI) for helpful discussions. We thank Matthew Burton and Eleanor Jones from the MCRI Flow Cytometry and Imaging Facility. This work was supported by the Stafford Fox Medical Research Foundation, L.E.W. Carty Trust, and the Novo Nordisk Foundation Center for Stem Cell Medicine (grant NNF21CC0073729).

## Author Contributions

R.B.W. conceptualised the project; D.L.T., H.B. and R.B.W. designed experiments; D.L.T., H.B., K.P., S.A., K.S., J.M. and R.B.W. performed experiments; D.L.T., L.G., M.N., S.S. and R.B.W. performed single-cell RNA-sequencing experiments; D.L.T., M.S., F.R. and R.B.W. performed bioinformatics analyses; S.L. provided influenza virus; E.N., A.E., M.R., F.R. and E.S. provided expert input on experimental design and data interpretation; R.B.W. and D.L.T. wrote the first draft of the manuscript. All authors critically reviewed and approved the final version of the manuscript.

## Conflict of Interest

F.J.R. receives institutional and salary support as a) a coinvestigator and subcontractor with the Peter MacCallum Cancer Centre for an investigator-initiated trial which receives funding support from Regeneron Pharmaceuticals; and b) a co-investigator on a translational research project funded by a Regeneron Pharmaceuticals grant.

---

**Supplemental Video 1. Relates to Figure 1.** Live cell confocal over 14 hours of iAT2 (marked by SFTPC-tdTomato) and iMacs (stained with CFSE).

**Supplemental Figure 1.**
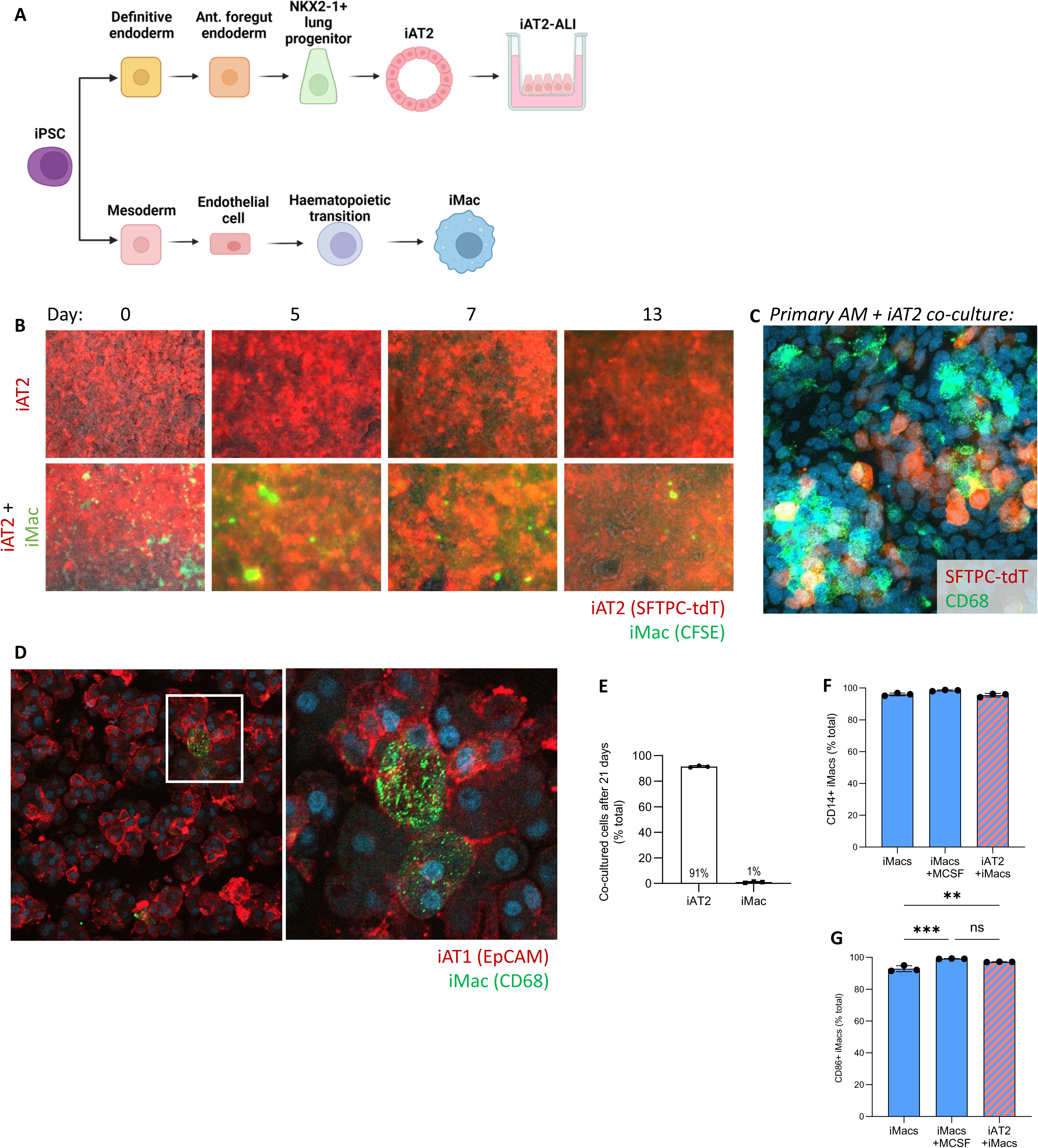
**Relates to Figure 1.** A) Schematic overview of iAT2 and iMac differentiation protocol. B) Live cell imaging of iAT2s (red, marked by SFTPC-tdTomato) and iMacs (green, labelled with CFSE) over 13 days. C) iAT2 (red, SFTPC-tdTomato) and alveolar macrophages (green, CD68) coculture with primary alveolar macrophages from the bronchoalveolar lavage. D) Immunofluorescence image of iAT1 (red, EpCAM) and iMac (green, CD68) coculture. E) Summary of flow cytometry data derived from day 21 cocultures days showing the relative proportions of iAT2s (NKX2.1-GFP+ SFTPC-tdTomato+) and iMacs (CD14+ CD45+). F) Flow cytometry analysis indicating the proportion of CD45+ cells expressing CD14 after 7 days cultured in CK-DCI, CK-DCI + M-CSF, or cocultured with iAT2s in CK-DCI. G) Flow cytometry analysis indicating the proportion of CD14+CD45+ cells expressing CD86 after 7 days cultured in CK-DCI, CK-DCI + M-CSF, or cocultured with iAT2s in CK-DCI.

**Supplemental Figure 2.**
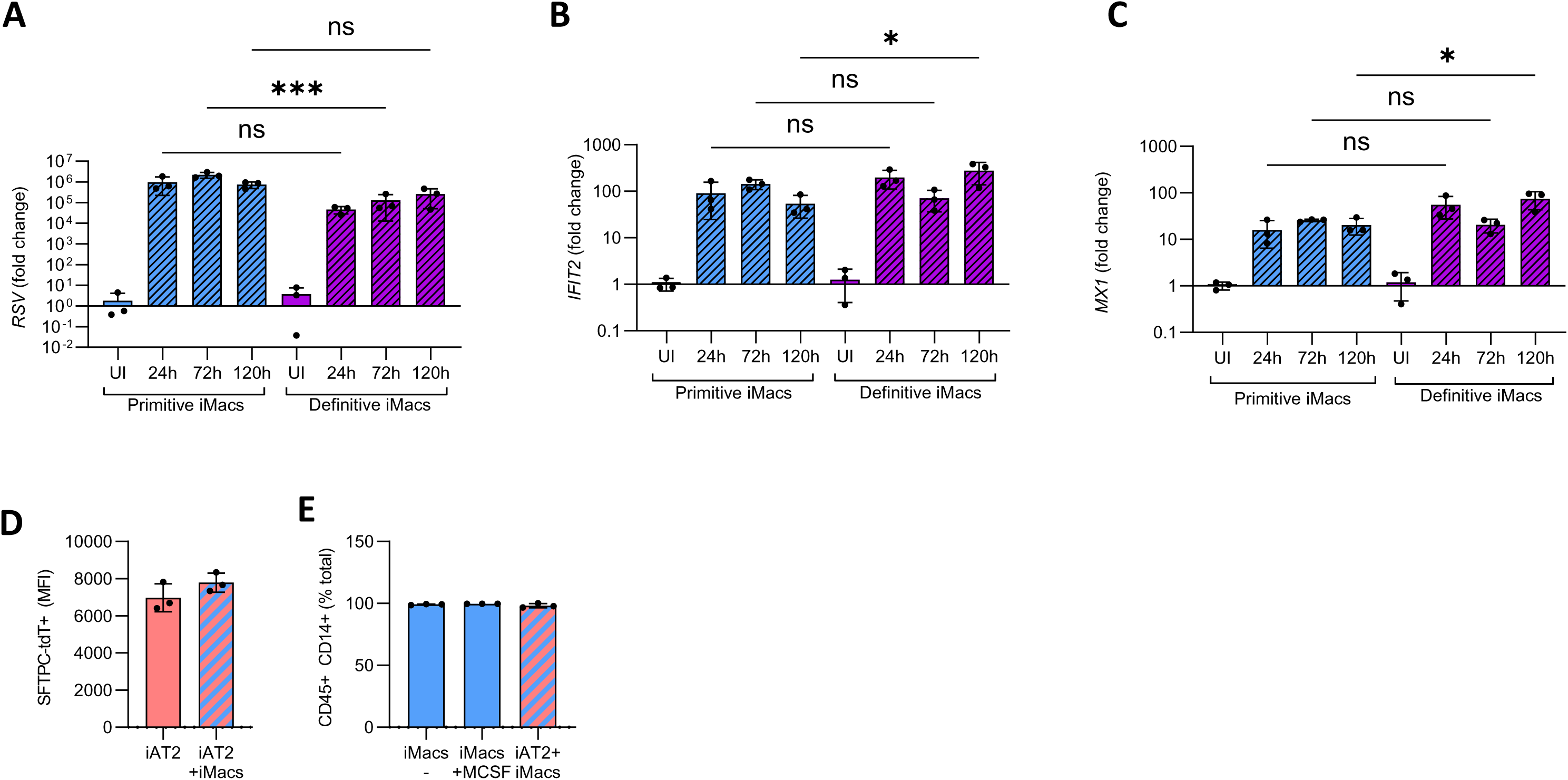
**Relates to Figure 1**. qRT-PCR analysis of experiments in which iMacs produced using primitive ^25^ or definitive ^38^ differentiation protocols were infected with RSV (MOI 1) for the times indicated. UI=uninfected. Cultures were analysed for expression of A) *RSV N,* B) *IFIT2* and C) *MX1*. D) iMacs produced using definitive differentiation protocol were cocultured with iAT2s at ALI. SFTPC-tdTomato expression measured by flow cytometry. E) Summary of flow cytometry experiments measuring the proportion of CD45+ cells expressing CD14+ (iMacs) following culture in CK-DCI, CK-DCI supplemented with M-CSF, or coculture with iAT2s. n = 3 experimental replicates of independent wells of a differentiation; error bars represent SD. Statistical significance was determined by an unpaired, two-tailed Student’s t test (two groups) or one-way ANOVA (>2 groups); *p < 0.05, **p < 0.005, ***p < 0.001.

**Supplemental Figure 3.**
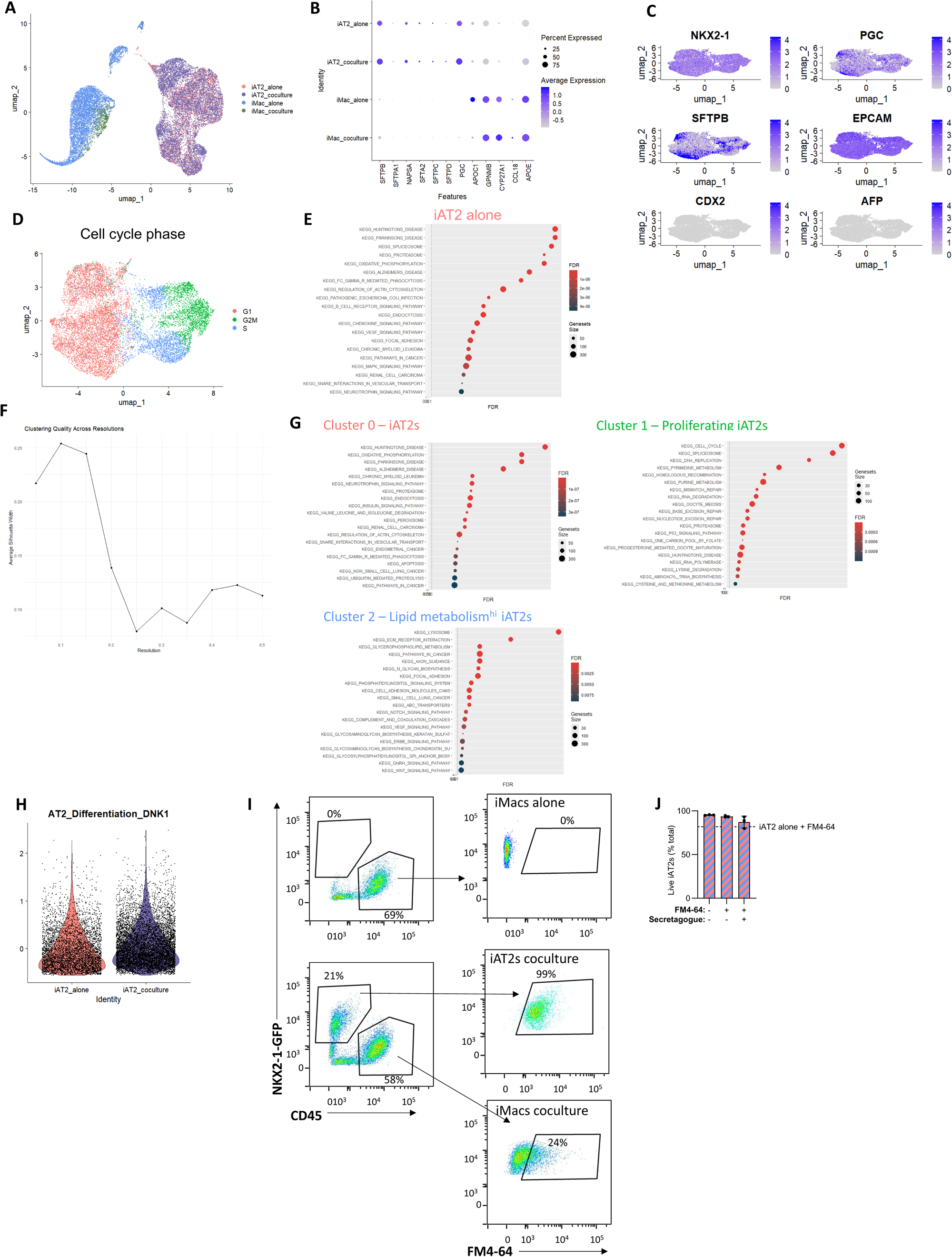
**Relates to Figure 2**. A) Uniform manifold projection (UMAP) of iAT2 alone (pink), iAT2 following coculture with iMacs (purple), iMacs alone (blue), or iMacs following coculture with iAT2s (green). B) Expression of markers associated with AT2s or macrophages in the adult human lung ^92^. C) UMAP of iAT2 markers (*NKX2-1, PGC, SFTPB, EPCAM*) and common non-lung endoderm lineages (*CDX2, AFP*) ^56^. D) UMAP of cell cycle phase of iAT2 alone and iAT2 following coculture with iMacs (Figure 2A). E) Gene set enrichment analysis of upregulated pathways in iAT2s alone. F) Silhouette width plot to determine clustering quality across Louvain clustering resolutions which determined a resolution of 0.1 was optimal. G) Gene set enrichment analysis of clusters identified at Louvain resolution 0.1. H) Module score of AT2 differentiation gene set ^56^. I) Flow cytometry gating for iAT2s and iMacs in the FM4-64 experiment (Figure 2I). J) Live iAT2s in FM4-64 experiment.

**Supplemental Figure 4.**
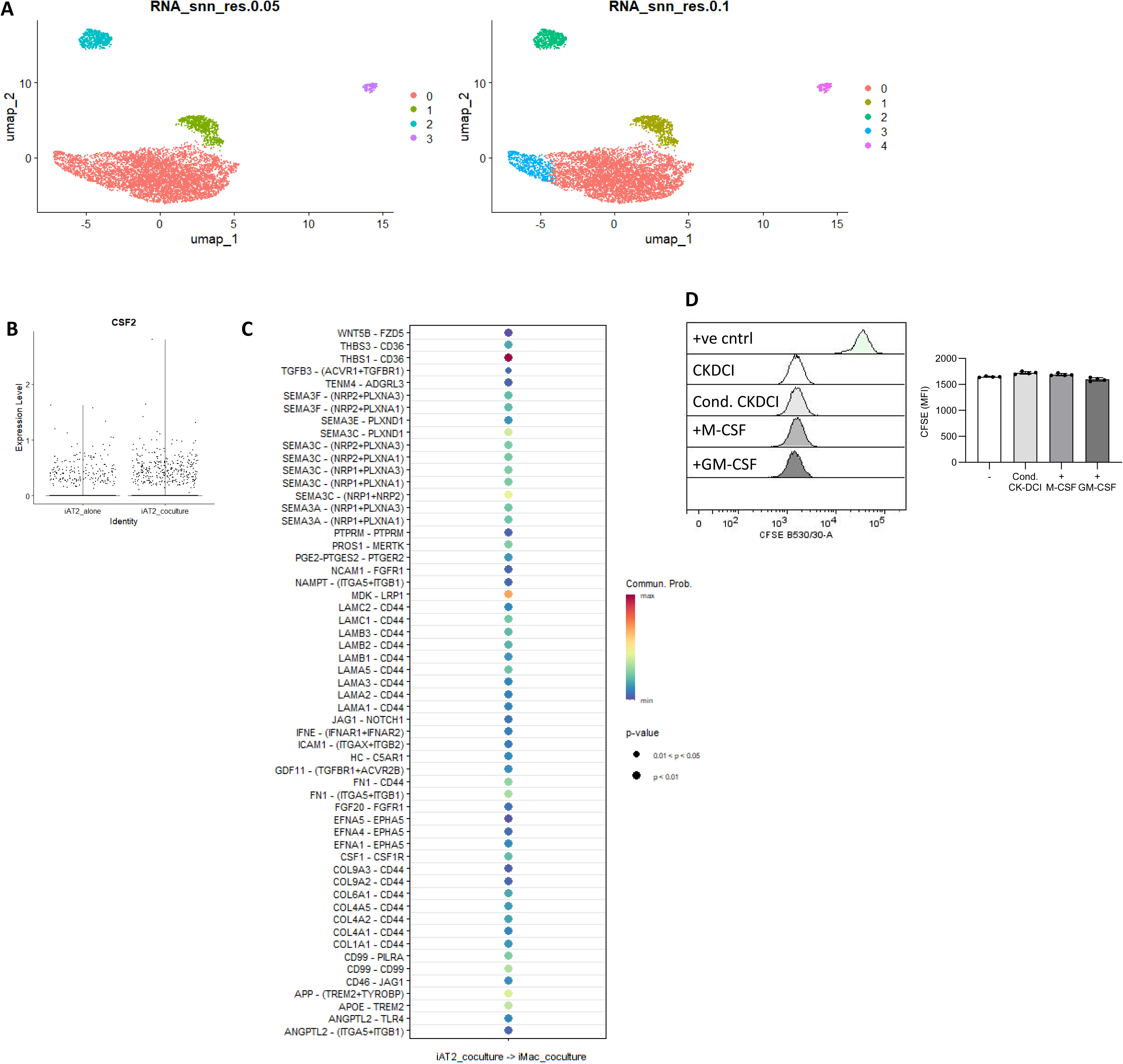
**Relates to Figure 3**. A) Uniform manifold projection (UMAP) of louvain clustering at a resolution of 0.05 and 0.1 of iMacs alone or iMacs following coculture (Figure 3A). B) *CSF2* expression in iAT2 alone (pink), or iAT2 following coculture with iMacs (purple) showing similar expression of *CSF2* expression between iAT2s alone and iAT2s in coculture. C) Ligand-receptors identified by CellChat analysis of iAT2s in coculture to iMacs in coculture. D) Proliferation of iMacs measured by CFSE dilution is unchanged by treatment with conditioned CK-DCI from iAT2s, or CK-DCI supplemented with M-CSF or GM-CSF for 72 hours.

**Supplemental Figure 5.**
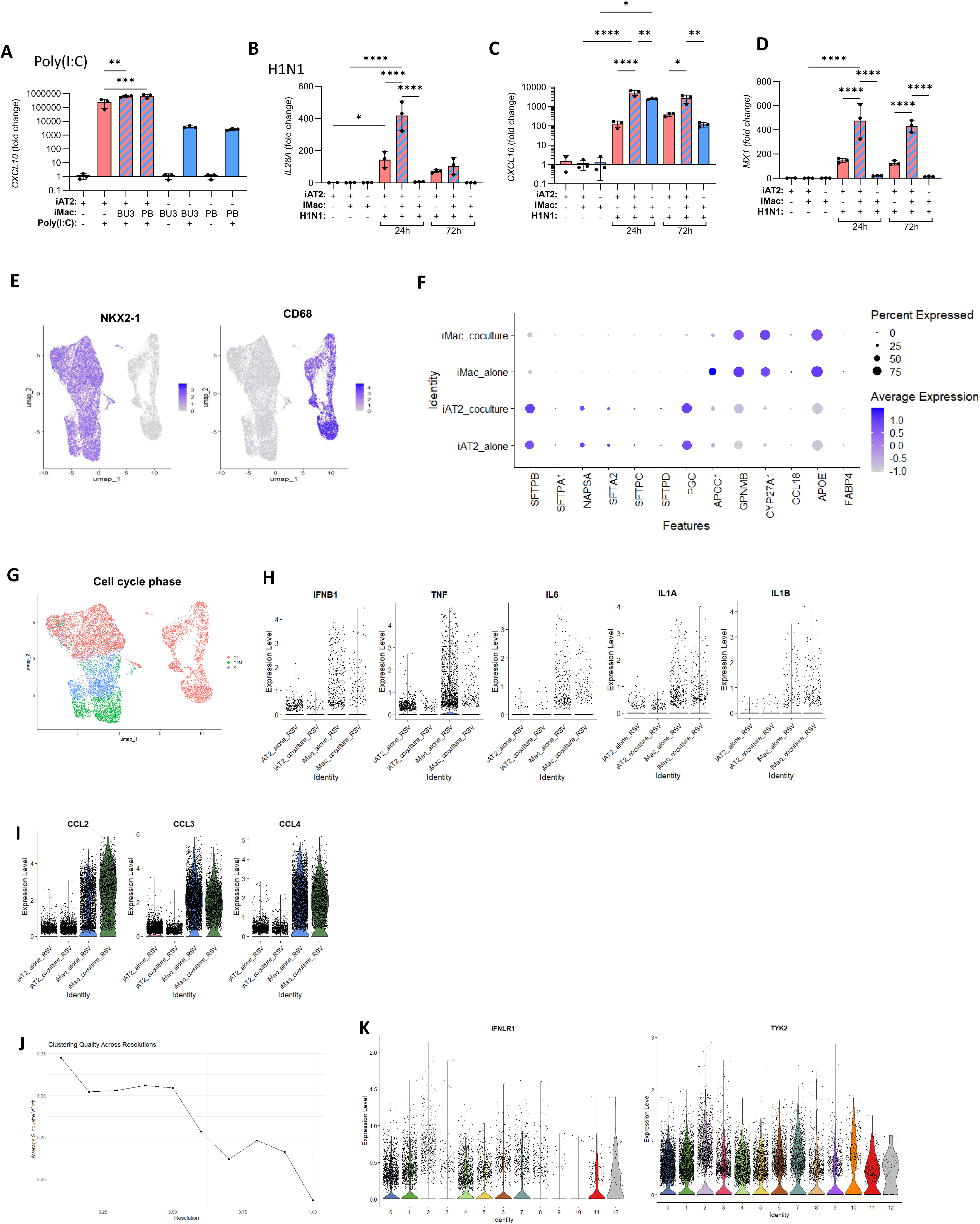
**Relates to Figure 4**. A) ALIs containing iAT2s alone or iAT2-iMac cocultures, or iMacs alone were stimulated with the TLR3 agonist, poly(I:C) for 24 hours. *CXCL10* expression measured by qRT-PCR. B) ALIs containing iAT2s alone or iAT2-iMac cocultures, or iMacs alone were infected with influenza A (H1N1 strain,, MOI 2) for 24 or 72 hours. *IL28A,* C) *CXCL10* or D) *MX1* were measured by qRT-PCR. E) NKX2-1 and CD68 expression in UMAP projection demarcating iAT2s (left) and iMacs (right) (Figure 4A). F) Expression of markers associated with AT2s or macrophages in the adult human lung ^92^. G) Cell cycle phase of iAT2s alone, iAT2-iMac cocultures or iMacs alone infected with RSV. H) Expression of antiviral cytokines and proinflammatory cytokines associated with macrophage expression during RSV infection. I) Expression of chemokines, primarily associated with macrophages, during RSV infection. J) Silhouette width plot to determine clustering quality across Louvain clustering resolutions which determined a resolution between 0.2-0.5 was optimal. K) IFNLR1 and TYK2 expression in clusters (Louvain resolution 0.4, Figure 4B).

**Supplemental Figure 6.**
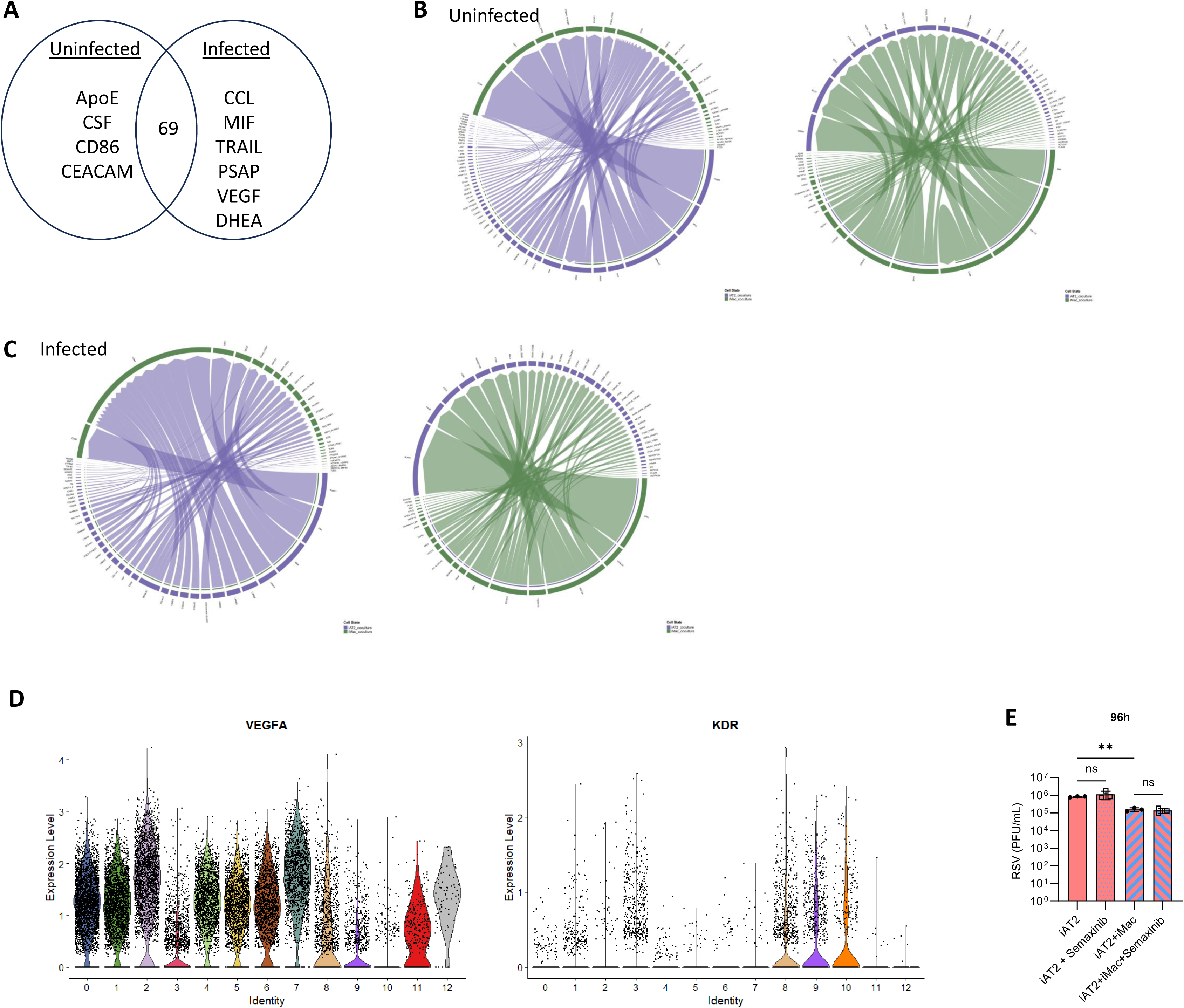
**Relates to Figure 5**. A) CellChat pathways between iAT2s and iMacs in cocultures common (69 shared pathways) or disparate between uninfected (ApoE, CSF, CD86, CEACAM) and RSV infected cultures (CCL, MIF, TRAIL, PSAP, VEGF, DHEA). C) Circos plot showing CellChat analysis between iAT2s (purple) and iMacs (green) in uninfected or C) RSV infected cocultures. D) VEGFA and KDR expression in clusters (Louvain resolution 0.4, Figure 4B). E) Shed RSV in the apical compartment at 96 hours of ALIs containing iAT2 alone or iAT2-iMac cocultures without or with semaxinib treatment was measured by plaque assay.

